# pathCLIP: Detection of Genes and Gene Relations from Biological Pathway Figures through Image-Text Contrastive Learning

**DOI:** 10.1101/2023.10.31.564859

**Authors:** Fei He, Kai Liu, Zhiyuan Yang, Yibo Chen, Richard D. Hammer, Dong Xu, Mihail Popescu

## Abstract

In biomedical literature, biological pathways are commonly described through a combination of images and text. These pathways contain valuable information, including genes and their relationships, which provide insight into biological mechanisms and precision medicine. Curating pathway information across the literature enables the integration of this information to build a comprehensive knowledge base. While some studies have extracted pathway information from images and text independently, they often overlook the correspondence between the two modalities. In this paper, we present a pathway figure curation system named pathCLIP for identifying genes and gene relations from pathway figures. Our key innovation is the use of an image-text contrastive learning model to learn coordinated embeddings of image snippets and text descriptions of genes and gene relations, thereby improving curation. Our validation results, using pathway figures from PubMed, showed that our multimodal model outperforms models using only a single modality. Additionally, our system effectively curates genes and gene relations from multiple literature sources. A case study on extracting pathway information from non-small cell lung cancer literature further demonstrates the usefulness of our curated pathway information in enhancing related pathways in the KEGG database.

## I. Introduction

MEDICAL literature has produced a large number of biological pathways in the form of images and texts at an incredible speed. These pathways usually describe the biological functional processes involving multiple genes and their interactions more intuitively and comprehensively provide abundant resources for the research of biology and the practice of precision medicine. For example, breast cancer (BC) is the most diagnosed malignant tumor in women, and the Hippo pathway is an evolutionary conserved biological signal pathway that plays a key regulatory role in various biological processes [1]. Therefore, in the fields of disease treatment and drug research, biologists and medical scientists need to devote themselves to the study of gene interaction in the latest articles, which plays a crucial role in guiding research and practice.

The extraction of pathway information and the construction of pathway databases remains challenging due to the rapid growth of publications and the scatter of pathway information in articles. Some public pathway databases such as KEGG [2], and Reactome [3] were built on manual curation. Although such handcrafted pathway databases are able to provide precise pathway knowledge, they are labor costly and time-consuming to mine and integrate genes and their interactions across thousands of medical articles. Some computational curation approaches were proposed to automatically mine genes and interactions from text and figures of literature. Researchers attempted to retrieve the gene and their interactions by text mining via Nature Language Processing (NLP) approaches. For example, PubTator Central [4] uses Named Entity Recognition (NER) as the core method to identify biological concepts from texts and is used to identify gene names in abstracts. Another example is to use the node2vec [5] model to extract different types of biological relationships from words, such as gene-gene relationships. However, they only extract information from the text, which inevitably leads to errors due to semantic complexity and noise interference, and their accuracy is limited.

While some researchers set their eyes on pathway diagrams taking advantage of computer vision technology. The extraction of pathway figures information in the literature is not as simple as extracting annotated regular pathway from KEGG, since the pathways from publications are figures that describe a disease or gene process according to the author’s research conclusions and certain biological rules, and they are regarded as individual images with no structured information at all. Hanspers, K. et al. extracted gene instances from pathway images by combining machine learning, Optical Character Recognition (OCR), and artificial curing [6]. Our previous work integrated the object detection model based on deep learning and Google Optical Character Recognition service to extract genes and their interactions from the pathways [7]. Although images express genes and their interactions in a more explicit way, the current image-mining-based approaches are still far from perfect performance. The current OCR tools and image analysis are not robust enough to unavoidable noises including diverse backgrounds, various fonts, irregular symbols, and rotated text from pathway figures, which challenged the detection of gene names and relations.

In summary, the performance of obtaining pathway information from the single modality of text or image hampered by their weakness, so we employed the contrastive learning method to overcome the weakness. Contrastive learning can incorporate with multiple modalities in a self-supervised way. It pulls together the modalities describing a same target and pushes apart the modalities denoting different targets at the embedding space. As a result, the learned discriminative embeddings can be adapted for the downstream tasks by small-scale fine-tuning. A variety of contrastive learning frameworks have been proposed, such as MoCo [8], SimCLR [9], BYOL [10], and SimSiam [11]. In the computer vision (CV) community, a representative contrastive learning work is Contrastive Language–Image Pre-training (CLIP) that connected text and image modalities to improve several typical CV tasks including image classification [12], object detection [13], image retrieval [14] and image generation [15].

Our main contributions can be summarized as follows:

- Upon the inspiration from CLIP, we proposed an image-text contrastive learning framework pathCLIP, for the recognition of genes and their relationships within pathway figures. This framework leverages the power of contrastive learning to enhance gene embeddings through the guidance of text embeddings.capitalization.
- Connecting this multi-modal model to a pathway identifier module and a pathway object detection module, we pipelined an automatic pathway curation system from pathway figures and evaluated for its effectiveness using an annotated dataset of pathway figures.
- Through a case study focused on non-small cell lung cancer, we demonstrated the capability of pathCLIP to unearth previously undiscovered pathways. It enriched the existing pathway knowledge available in databases like KEGG.
- We published our code and annotated datasets to improve reproducibility, which was beneficial to a wide range of researchers.

## II. Method

### A. Data Preparation

We collected 21,273 publications from PubMed database using the query keywords signal pathway according to their accessibility. To enable downstream analysis, we used PyMuPDF [16] to extract embedded figures, text, and the structural layout of the text within each publication. We manually checked each figure and distinguished 5,693 biological pathway figures from all non-pathway figures in the publications. These pathway figures, as well as non-pathway figures, served as positive and negative samples for building the pathway identifier. To annotate the text, arrows, and T-bars within the pathway figures (representing genes and gene relations), we employed the labelme [17] tool. This annotation involved creating bounding boxes around objects and categorizing them for use in object detection modeling.

Then we created two image-text multi-modal datasets, i.e., Gene Detection Dataset (GD Set) and Gene-Relation Extraction Dataset (GRE Set). We started from all labeled genes in pathway figures to retrieve all their mentioning sentences from their source publications. All these sentences were paired the cropped gene slices from the pathway figures to define an image-text multi-modal gene description set for contrastive learning.

For gene-relation extraction, we searched for sentences describing annotated gene relations in the pathway figures. This involved examining image captions, cited paragraphs, and other relevant sources. We identified 1,904 sentences covering any two genes involved in annotated gene relations. These sentences were used for fine-tuning BioBERT on relation extraction tasks. Additionally, these sentences were paired with cropped gene-relation images for contrastive learning.

Detailed information about the data was provided in Table I.

**TABLE I.** Collected data descriptions

| Data level | Task | Source and label | Division |
| --- | --- | --- | --- |
| Figure | Pathway figure identifier | Figures obtained from publications manually labeled as pathways and non-pathways. | Training data: 400 figures<br>The positive and negative samples are roughly 1:1. And do five-fold cross-validation on 193 negative samples and 30 positive samples. |
| Object | Object detection | Arrows/T-bar lines, heads, gene text and gene-relation triples bounding boxes in annotated pathway figures are labeled by the 'labelme' tool. | Training data: 4,040 pathway figures<br>Independent testing data: 45 pathway figures<br>There are 34,127 arrows, 30,245 T-bars, 89,740 genes in training data and 561 genes and 367 gene relations in independent testing data. |
| Sentence | BioBERT fine-tuning on RE | Sentences in publications describing gene activation relation is marked as "1" and inhibition relation is marked as 0. | Training data: 1,277 sentences<br>Testing data: 677 sentences<br>The ratio of activation to inhibition is approximately 1: 1. |
| Image-text pair | Gene detection by pathCLIP (GD set) | Sentences from literature were paired the cropped gene slices from the pathway figure, which cover 25 types of genes. | Training data: 6,313 image-text pairs<br>Testing data: 1,939 text<br>The proportion of training and testing samples is roughly 4: 1. |
|  | Gene-relation extraction by pathCLIP (GRE set) | Sentences describing gene relations and their cropped gene-relation slices from the pathway figures were paired, which have two categories: activation and inhibition. | Training data: 3,658 image-text pairs.<br>Testing data: 865 text |

### B. Overview of Pipeline

We developed pathCLIP, a comprehensive framework designed to extract genes and their relations from biological pathway figures and associated text descriptions. Fig. 1 illustrates the overview of pathCLIP, which mainly includes three key components: (1) the image module on the left to extract visual embedding from pathway figures, (2) the text module on the right to generate text embedding from article text, and (3) the image-text contrastive learning module in the middle to achieve the tasks of gene entity recognition and relation extraction.

**Fig. 1.**
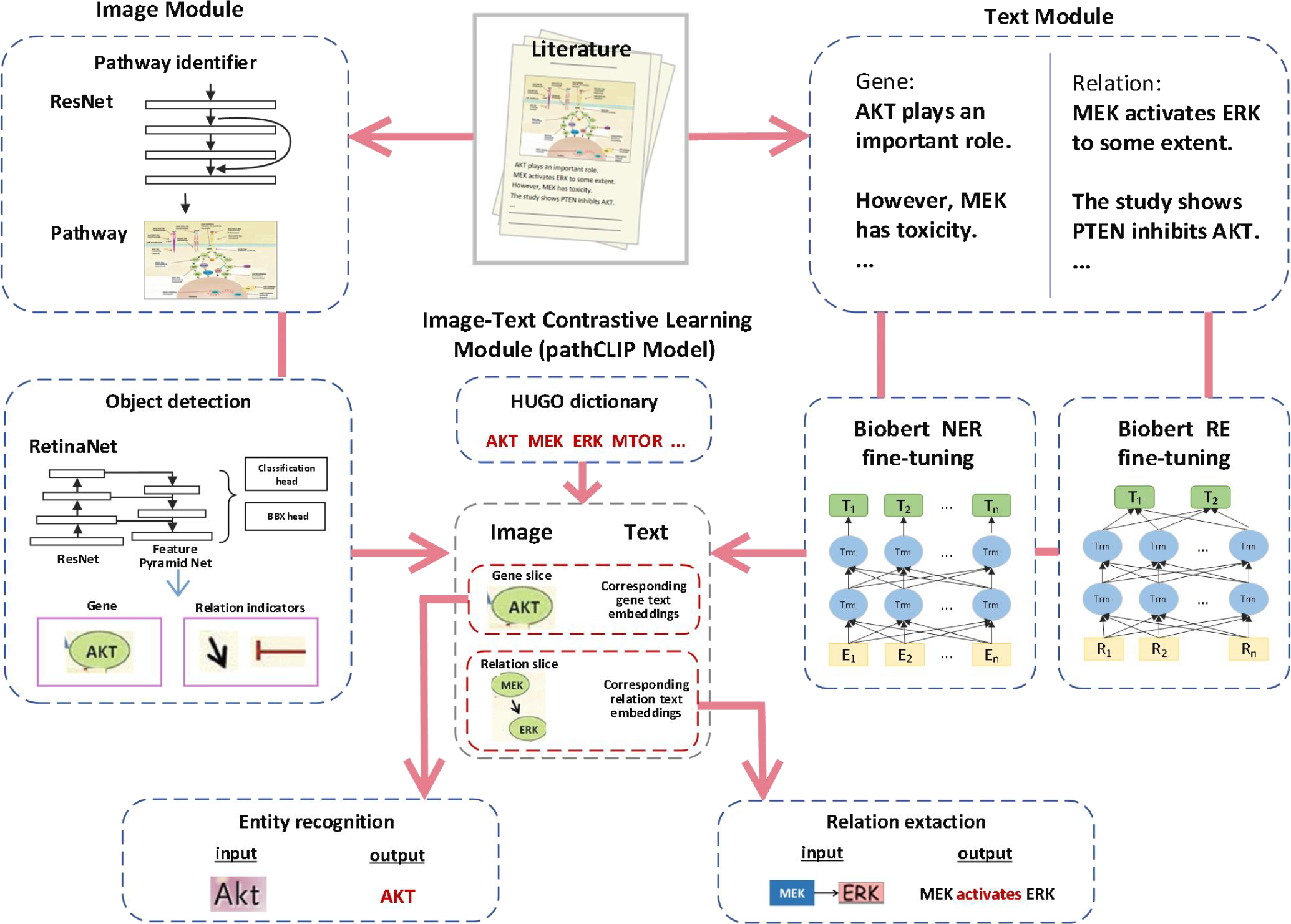
Proposed Pipeline Overview. This figure provides an overview of our pipeline, consisting of three key modules. On the left, the image module extracts visual embeddings from pathway figures. On the right, the text module generates text embeddings from the article text. In the middle, our image-text contrastive learning module (pathCLIP) is shown, which facilitates gene entity recognition and relation extraction tasks.

### C. Pathway Figure Identifier

To facilitate the automatic curation pipeline, it is essential to differentiate pathway figures from various other image types commonly found in the biomedical literature, such as flow charts, organ maps, and chemical molecular formula images. Our first step is to distinguish the pathway figure from other images. We fine-tuned a pretrained ResNet50 [18] on ImageNet with our labeled images from the literature. ResNet50 stacks 50 convolution layers with skip connections to capture the multi-scale and abstract features from images. The skip connections were instrumental in mitigating the vanishing gradient issue often encountered in deep architecture. Given the binary prediction objective (pathway or non-pathway), the binary Cross-Entropy Function [19] was applied as the loss function for training the Resnet50. Adam [20] was adopted as the optimizer, and the learning rate was set to 1e^-5^. To avoid possible bias caused by imbalanced training data, we randomly extracted the same number of non-pathway figures with pathway figures from the full training set for each training epoch. The training process continued until reaching a maximum of 300 epochs. At this point, the model had effectively learned to classify and filter out irrelevant figures, thereby serving as a reliable pathway figure identifier for downstream analysis. In order to fairly evaluate the model, we also conducted five-fold cross validation in the evaluation stage. The network structure, and process flow chart of Pathway identifier are shown in the Fig. 2(a).

**Fig. 2.**
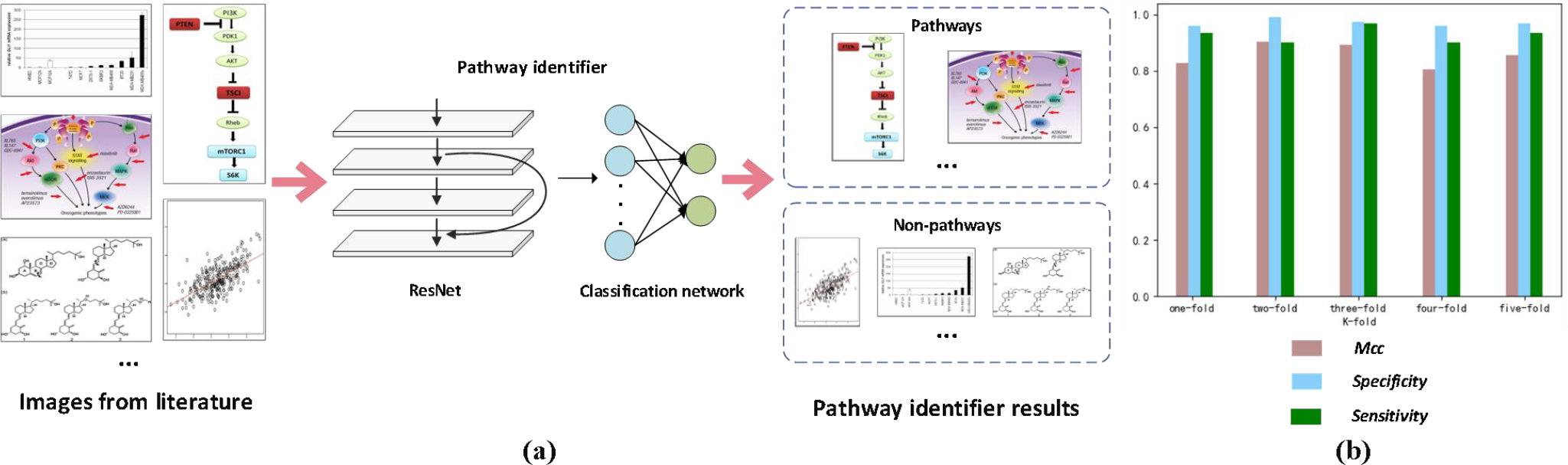
Pathway Identifier. (a) illustrates the figure retrieval steps and the architecture of the Pathway Identifier. The input image, extracted from the literature, is passed through the Pathway Identifier, comprising ResNet and a Classification network. The output indicates whether the image belongs to the pathway or non-pathway category. (b) presents the performance of the Pathway Identifier, measured in terms of precision, recall, and F1-score.

### D. Object Detection from Pathway Figures

To mine the symbols defining genes and relationships from pathway figures, we trained an object detector using RetinaNet architecture [21] to identify and localize text, arrows, and T-bars, which respectively represent genes, activation relations and inhibit relations. With the locations of these symbols on pathway figures, we can infer the genes and their relations on the pathway figures.

Our RetinaNet employed pretrained ResNet101 [18] on ImageNet as the backbone network to provide multi-scale visual feature maps. A series of anchors were generated on each scale feature map according to the size and aspect ratio as 0.5, 1 and 2 in our practice, providing region proposals for searching objects. The IOU (Intersection Over Union) [22] was used to determine if any objects fell into the anchor. Anchors with IOU greater than 0.6 were marked as positive, while anchors with IOU less than 0.4 were considered negative. anchors with IOU between 0.4 and 0.6 were ignored to reduce the computation. Following the multi-scale feature maps, RetinaNet appended two subnets to predict the category and coordinates of each anchor point on the input images. The class subnet mapped each anchor’s feature map to a score through a softmax function, while the regression subnet processes the same feature maps into 4-dimensional coordinates forming a bounding box. In the classification subnet, the Focal loss [21] was utilized to address the unbalance between positive and negative cases.

In the regression subnet, we use a smoothed L1 loss function [23] to measure the difference between the anchors and predicted bounding boxes. These two subnets were optimized by Nadam optimizer [24] with a learning rate starting from 5e^-4^ and decaying to 1e^-5^. Once the RetinaNet completed its training, the categories and locations of arrows/T-bars and arrowheads/T-bar heads from any input pathway images were detected by the trained RetinaNet. In inference, a 0.7 confidence threshold and non-maximum suppression operation [25] were used to filter out low confidence and duplicate boxes on the same object. Considering that arrows and T-bar lines denote activation and inhibit interactions and their heads denote the direction of the interaction, we trained two RetinaNets to detect arrows/T-bar lines and their heads separately using annotated relation snippets from collected pathway figures. The network structure, process of Pathway object detection is shown in the Fig. 3(a).

**Fig. 3.**
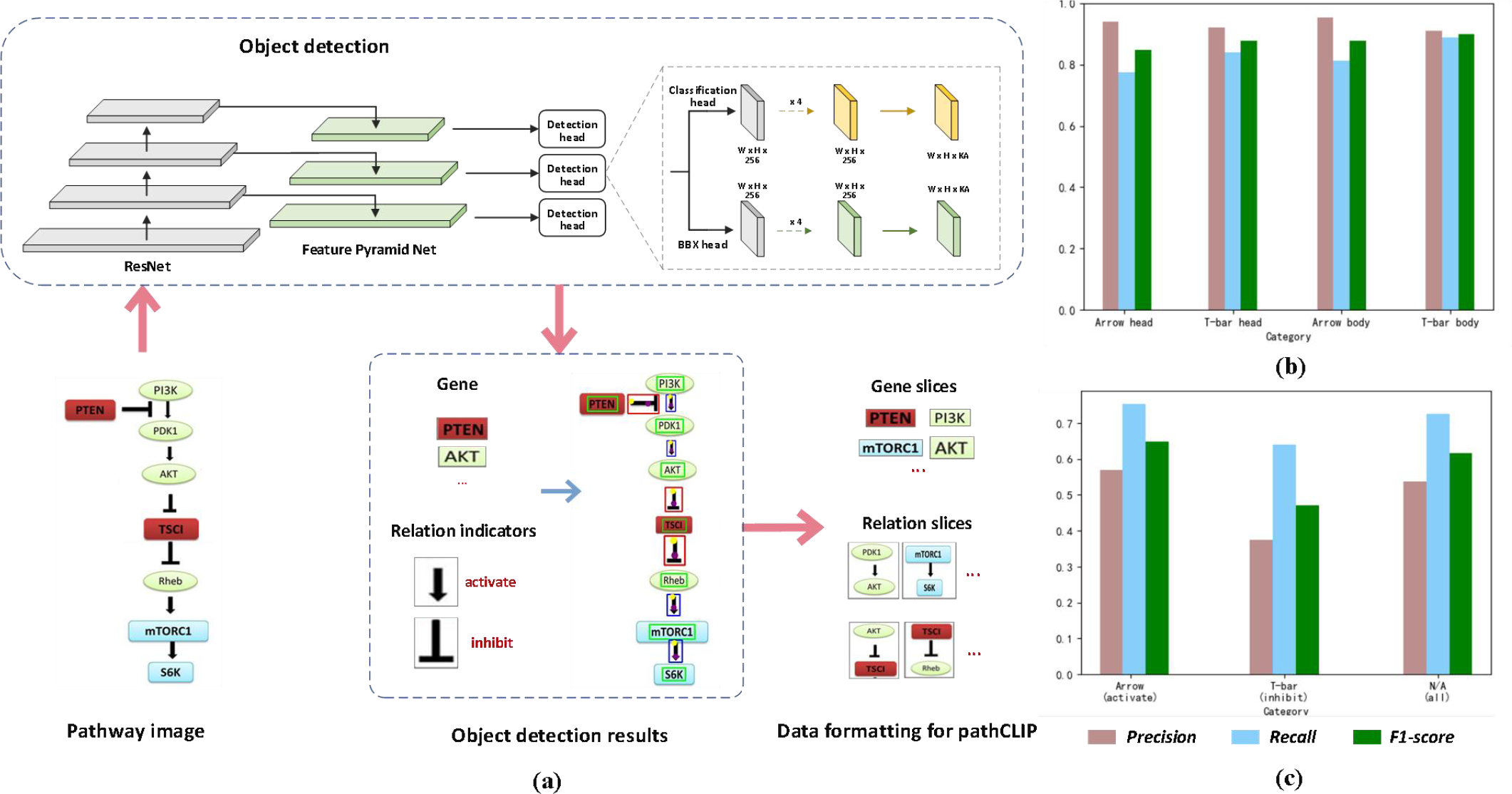
Pathway Object Detection. (a) illustrates the image data preprocessing steps and the architecture of the object detection model. The pathway image is input into the object detection model, which identifies text boxes, arrows, and T-bars. These elements are then organized into image data acceptable for CLIP. (b) Performance Metrics for Arrow and T-Bar Detection. (c) Performance Metrics for Gene-Interaction Recognition Detection

### E. BioBERT Fine-tuning on tasks of NER and RE

As valuable sources for gene and relation mining, text descriptions provide clear representations essential for curation. We conducted fine-tuning operations using BioBERT [26], a biomedical text mining variant of BERT [27] pretrained on biomedical corpora. This fine-tuning process aimed to enhance the adaptability of BERT to biomedical text, ultimately yielding text embeddings for genes and gene relations for image-text contrastive learning.

BioBERT is a biomedical specialized version of BERT, which stacks multiple transformer blocks [28]. Each block includes multi-head attention layers and a LayerNorm layer [29], enabling it to capture contextual information and dependencies between input words effectively. For gene text embedding, we directly used a fine-tuned BioBERT for Named Entity Recognition (NER) tasks using the BC2GM corpus provided by HuggingFace [30]. This fine-tuned model, named BioBERT-NER, facilitated superior embedding of input sentences to represent gene representation.

To obtain embeddings for gene relations, we further fine-tuned BioBERT using our annotated sentences that describe gene relations. Based on the two types of gene relations outlined in Table I, we performed fine-tuning in a binary classification setting with BioBERT-RE. During training, we utilized the binary cross-entropy loss function [19], with optimization carried out using AdamW [31] and a learning rate set to 5e^-5^ over 300 epochs to ensure model convergence. This refinement of BioBERT empowered it to generate text embeddings representing potential gene-disease relations from input sentences, and provided a good start point for further training a model extracting our gene-gene relationships. We presented some annotated sentences at the gene and gene-relation levels in Fig. 4(a). Additionally, a schematic diagram illustrating the BioBERT model’s involvement in NER and RE downstream tasks also was depicted in Fig. 4(b).

**Fig. 4.**
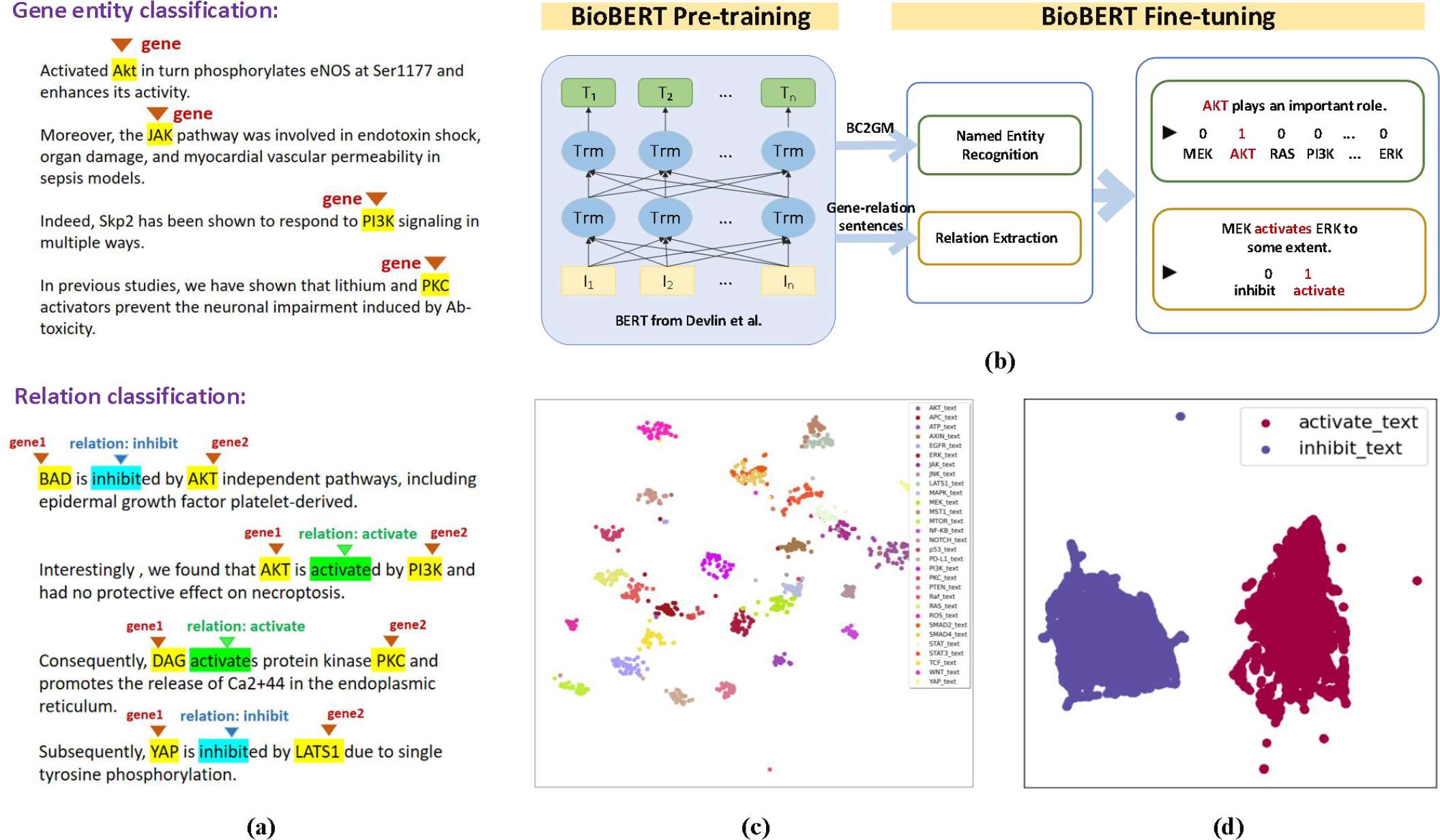
Text BioBERT Fine-Tuning Details. (a) Structure of text data used for gene entity detection and relation extraction tasks, highlighting key information related to genes and relations. (b) BioBERT Model Pre-Training and Fine-Tuning is depicted. These models are employed to extract text features from sentences describing genes and relations, respectively. (c) Embedding Distribution of BioBERT-NER Model on Gene Entity Detection Texts. (d) Embedding Distribution of BioBERT-RE Model on Relation Extraction Texts.

### F. Contrastive Learning on Pathway Figures and their Text Descriptions

Inspired by OpenAI’s image-text contrastive learning framework CLIP, we extended its capabilities to biomedical pathway figures and associated publications. CLIP learns visual features guided by natural language supervision, fostering a deep understanding of the relationship between images and their textual descriptions. We leveraged this concept to enhance the visual representations of genes and gene relations within pathway figures using textual supervision.

Our contrastive learning approach encompassed two distinct tasks: gene recognition and gene-relation extraction. For gene recognition, we utilized the fine-tuned BioBERT-RE as the text encoder and employed a pre-trained ResNet50 from ImageNet as the image encoder. CLIP operates within a shared embedding space, where semantically related image-text pairs are positioned closely to each other. To achieve this, we incorporated two linear projectors following the text and image encoders. These projectors mapped the generated embeddings to a unified 256-dimensional space, as depicted by (1) and (2):

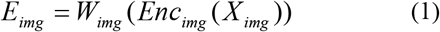

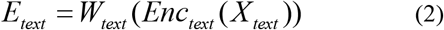

The image (X_img_) and text (X_text_) in a batch are expressed using their respective encoders, and then mapped to the shared 256-dimensional space using linear projector W_img_ and W_text_. The objective of our contrastive learning was to push similar image-text pairs (positives) closer in the shared embedding space while pushing dissimilar pairs (negatives) apart. To reach this goal, we created input batches consisting of *N* gene snippets from our annotated pathway figures and its corresponding mentioning sentences. Hence, there would be *N* positive pairs and *N*^*2*^ *− N* negative pairings in each batch. A symmetric cross entropy loss was used to train the encoders and projectors by maximizing the cosine similarity of embeddings of positive pairs while minimizing the cosine similarity of the embeddings of negative ones. The model was trained using backpropagation and Adam optimizer [20] with learning rate 1e^-4^ for image encoder, 1e^-5^ for text encoder and convergence criteria. The algorithm of contrastive learning is shown as follows.

#### Algorithm 1 Pseudo Code of Contrastive Learning

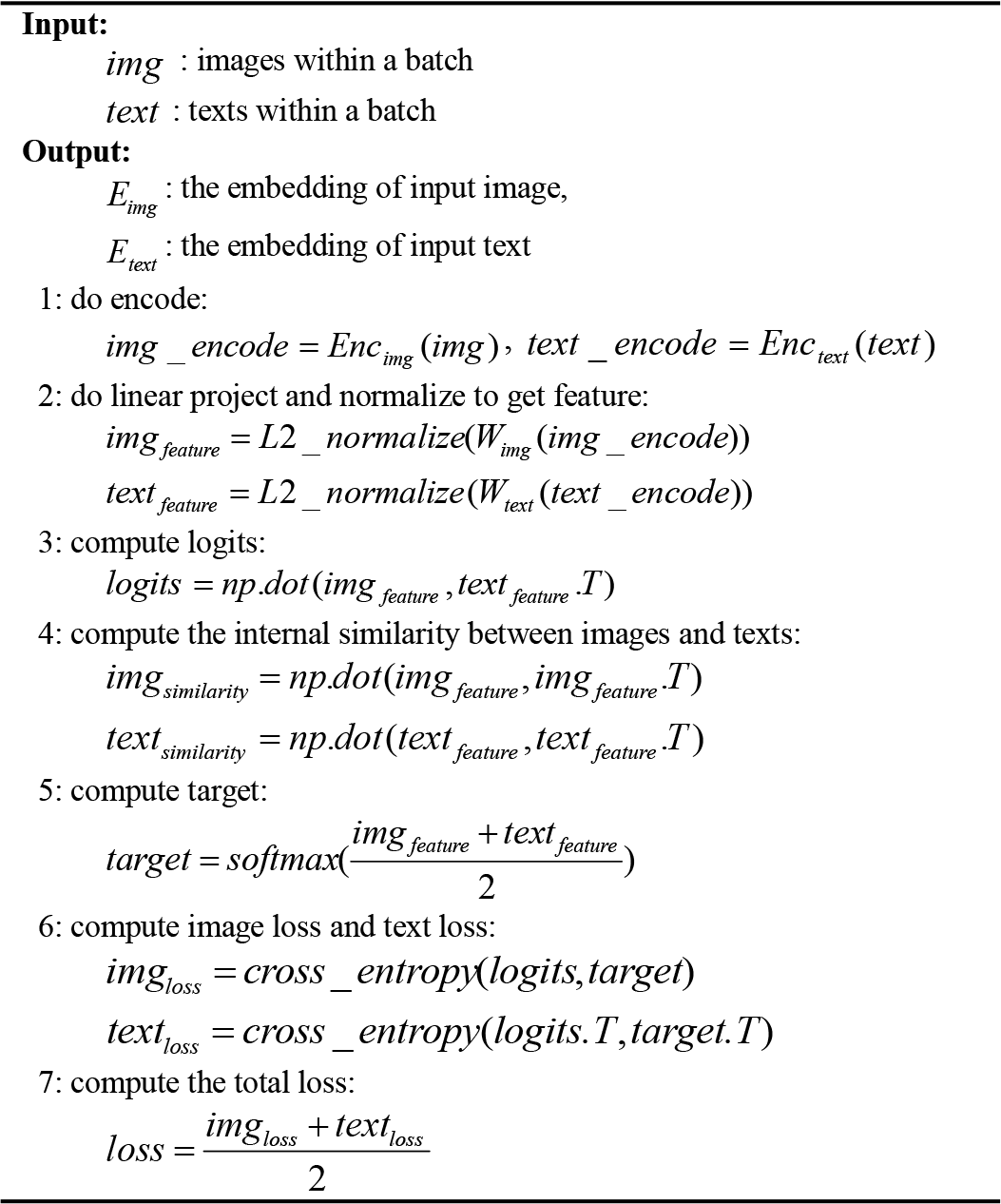

Following the contrastive learning phase, pathCLIP was equipped to perform gene name recognition by processing an unseen cropped image containing a gene from a pathway figure along with a prompt text. Given our current focus on human genes, we employed a specific image description prompt, *‘a snippet of [placeholder]’* and combined it with all possible gene names sourced from HUGO [32] at [placeholder] to create a set of candidate prompts. Next, pathCLIP measured the cosine similarity between the embedding generated from the cropped image and each candidate prompt. Higher similarity scores indicated a better match between the image and a specific prompt. Consequently, the prompt that exhibited the highest similarity with the image among all candidates was retrieved. The gene name contained within the retrieved prompt was predicted as the recognized gene name for the input cropped image. In this manner, pathCLIP efficiently identifies and assigns gene names to images based on their textual descriptions. The training of contrastive learning and inference process for gene name recognition are shown in Fig. 5(a).

**Fig. 5.**
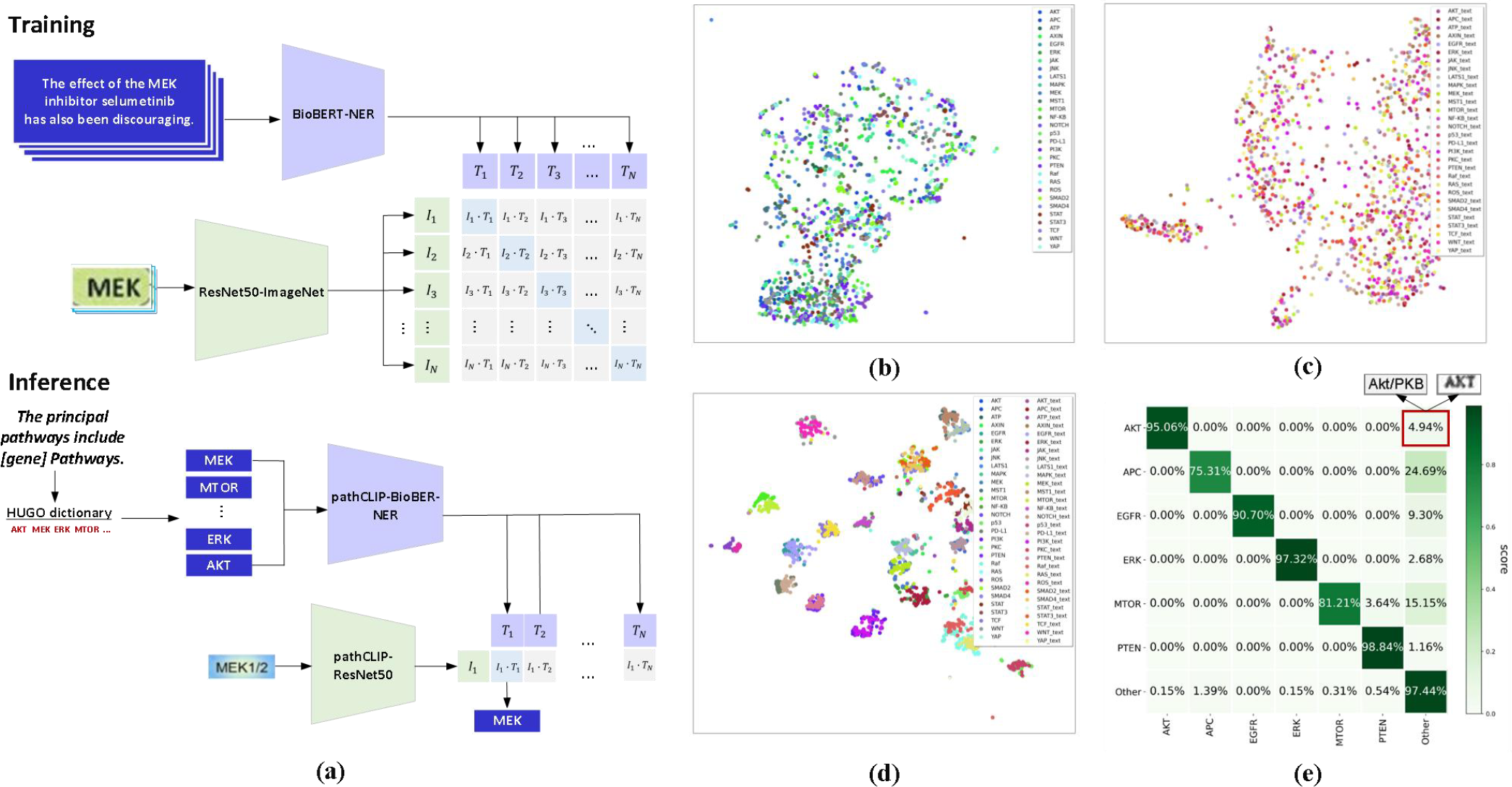
pathCLIP for Gene Entity Detection. (a) The training and inference processes of pathCLIP and its integration with CLIP. (b) Embedding Distribution of ResNet50 on Entity Detection Images. (c) Embedding Distribution of BioBERT on Entity Detection Texts. (d) Embedding Distributions of pathCLIP on Entity Detection Images and Text. (e) Confusion Matrix for Gene Entity Detection by pathCLIP.

In the context of gene-relation extraction, we employed a similar contrastive learning approach as before and utilized the fine-tuned BioBERT-NER model as our text encoder. Our training data consisted of gene-relation snippets extracted from annotated pathway figures and their corresponding text sentences. The same training settings were applied for contrastive learning in the gene-relation extraction task. Subsequently, we designed a parallel inference strategy for relation extraction.

For relation extraction, our prompt description took the form *‘gene1 [placeholder] gene2’* where gene1 and gene2 represented the recognized gene names obtained from the gene snippets extracted from the cropped relation images using the previously mentioned contrastive module. The [placeholder] was then replaced with either ‘activates’ or ‘inhibits’ to form two candidate prompts for inference. To determine the predicted gene relation for a given input cropped relation image, we sent both the unseen cropped relation images and all candidate prompts to the trained pathCLIP model. The model compared their embedding similarities and selected the best-matched prompt as the predicted gene relation for the input cropped relation image. The training of contrastive learning and inference process for gene name recognition are shown in Fig. 6(a).

**Fig. 6.**
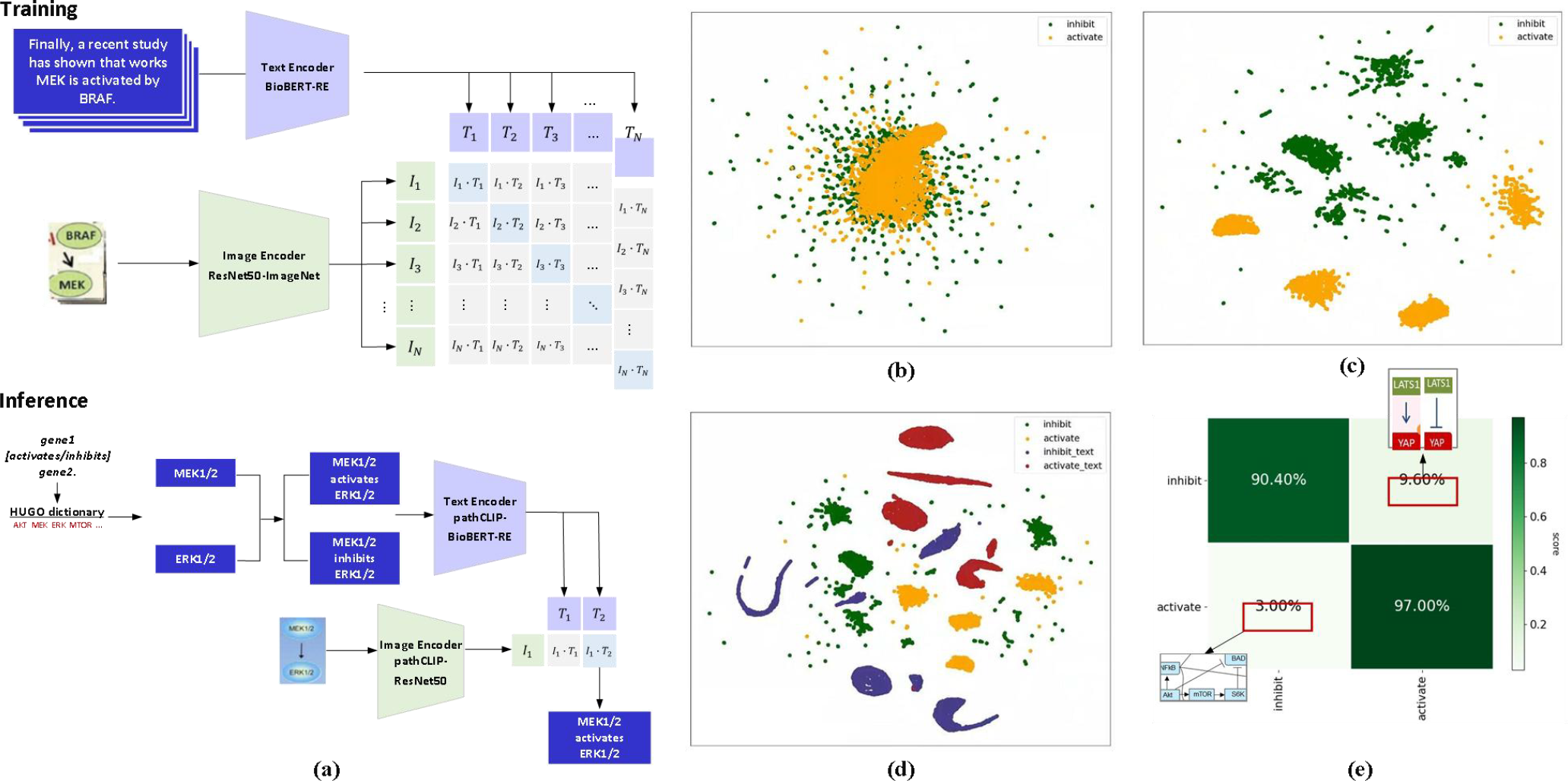
pathCLIP for Relation Extraction. (a) The training and prediction processes of pathCLIP and its integration with CLIP for relation extraction. (b) Embedding Distribution of ResNet50 on Relation Extraction Images. (c) Embedding Distribution of pathCLIP-ResNet50 on Relation Extraction Images. (d) Embedding Distributions of pathCLIP on Relation Extraction Images and Text. (e) Confusion Matrix for Relation Extraction by pathCLIP.

### G. Evaluation Metrics

As illustrated in (3)-(5), the performance metrics employed to assess the contrastive model included precision (the probability of correctly identifying a positive sample among all the samples predicted as positive), recall (the probability of correctly identifying a positive sample among all the actual positive samples), and F1-score (a composite metric that balances both precision and recall, ensuring they achieve their highest levels simultaneously). Specifically, TP (True Positive) is the true positive rate, FP (False Positive) represents the false positive rate, TN denotes the true negative rate. and FN (False Negative) is the false negative rate.

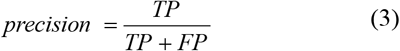

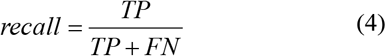

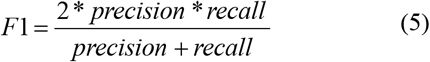

As depicted in (6)-(8), the metrics utilized for evaluating the pathway identifier model covered MCC (Matthews Correlation Coefficient), specificity (true negative rate), and sensitivity.

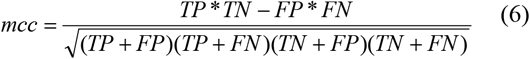

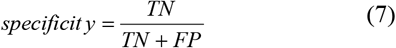

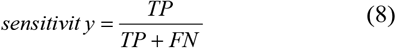

## III. Results

### A. The Performance of Pathway Identifier and Object Detection

First of all, we reported the performance evaluation of our pathway identifier using a 5-fold cross-validation approach, as summarized in Table II. We assessed the model’s discriminative power for distinguishing pathway figures from non-pathway figures by measuring the MCC, Specificity, and Sensitivity.

**TABLE II.**
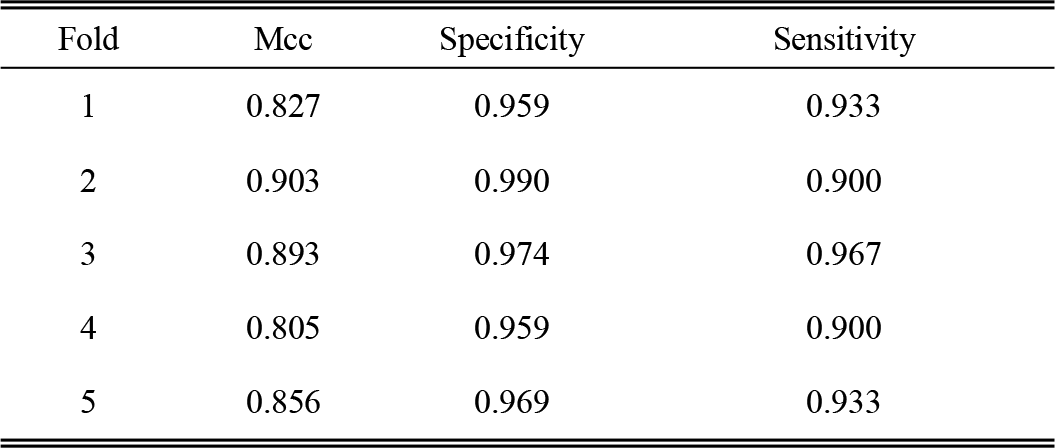
Performance of pathway identifier

| Fold | Mcc | Specificity | Sensitivity |
| --- | --- | --- | --- |
| 1 | 0.827 | 0.959 | 0.933 |
| 2 | 0.903 | 0.990 | 0.900 |
| 3 | 0.893 | 0.974 | 0.967 |
| 4 | 0.805 | 0.959 | 0.900 |
| 5 | 0.856 | 0.969 | 0.933 |

Our results demonstrated that pathCLIP consistently achieved impressive performance across these metrics. To further validate our model’s robustness, we conducted an evaluation on an independent test set, comprising 193 negative samples and 30 positive samples, the results of which are depicted in Fig. 2(b). Surprisingly, despite the inherent class imbalance between positive and negative samples, our training strategy, which employed balanced sampling, ensured that our model did not exhibit any biased predictions during the independent test. Upon reviewing instances where our model failed, we found that most false positives were falling into diagrams of methodology flowcharts, which were visually similar to pathway figures to some extent. Nevertheless, the pathway identifier established a robust foundation for subsequent stages of our study.

Fig. 3(b) and Fig. 3(c) illustrated the object detection performance in terms of precision, recall, and F1-score for arrows, T-bars, and gene interactions. These results aligned with our previous work [7], reinforcing the groundwork for the subsequent application of contrastive learning in gene and gene-relation identification.

### B. The enhanced embedding from Fine-tuning BioBERT

To explore the distinctiveness introduced by fine-tuning BioBERT, we visualized the embeddings generated by BioBERT without fine-tuning and those obtained from fine-tuned BioBERT (named BioBERT-NER). We leveraged the UMAP technique [33] to project these embeddings into a 2-dimensional space, allowing us to visualize the relationships among sentences describing genes. Fig. 4(c) presented the results of projecting BioBERT-NER embeddings. Notably, in this embedding space, sentences that described the same gene tend to be closer to each other than to sentences discussing different genes. This suggested that the embeddings generated by BioBERT-NER have a propensity to capture the essence of genes mentioned in input sentences.

Concurrently, we applied a similar procedure to project the embeddings produced by the fine-tuned BioBERT-RE model into a 2-dimensional space, visualizing the distribution of sentences describing gene relations in Fig. 4(d). Here, it is evident that sentences elucidating the same activate/inhibit gene relations tend to form distinct clusters, demonstrating a higher degree of coherence compared to randomly selected sentences.

### C. Gene Recognition by pathCLIP

To illustrate the effectiveness of pathCLIP, we conducted performance comparison involving the OCR approach presented in our previous work [7] and pathCLIP in the context of gene recognition on the detected gene image slices, with results summarized in Table III. Notably, pathCLIP delivered significant improvements in terms of precision, recall, and F1 score. These enhancements might be beneficial from the multi-modal contrastive learning process, where the image encoder in CLIP emerged the discriminative power learned from the text encoder’s supervision. As a result, the image features were enhanced, leading to superior gene recognition outcomes.

**TABLE III.**
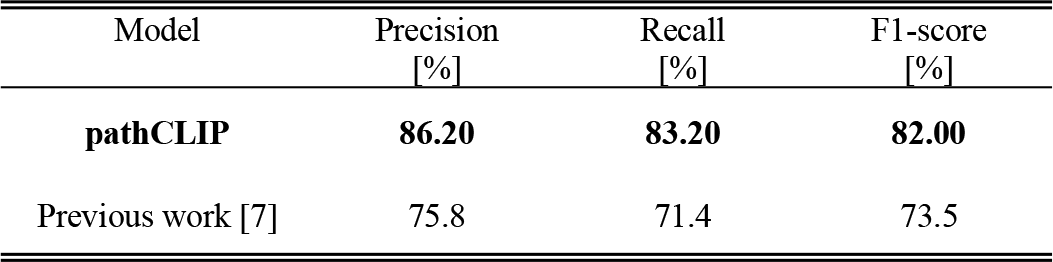
Gene recognition performance of pathCLIP on detected gene image slices

To provide further insight into the distinctiveness of image features, we conducted UMAP visualizations for both fine-tuned ResNet50 and ResNet50 from pathCLIP, as shown in Fig. 5(b) and Fig. 5(d), respectively. Fig. 5(b) depicted the image features from fine-tuned ResNet50, where the points appear scattered, lacking a clear clustering pattern. In contrast, Fig. 5(d) showcased the image features generated by pathCLIP, where they naturally cluster according to their associated gene names. This observation highlighted the limitations of ResNet50’s image features, which, without textual guidance, struggle to handle image resolutions and noise, resulting in reduced distinctiveness.

To complement our analysis, we examined the distribution of text features extracted from the pre-trained BioBERT model, as depicted in Fig. 5(c). In this generic large language model, no evident clustering areas for each gene category were observed. This suggested that BioBERT, in its pre-trained form, required further fine-tuning to adapt its text feature generation towards gene names.

We further explored the synergy between image and text encoders in pathCLIP by visualizing their embeddings after contrastive learning in Fig. 5(d). These embeddings demonstrated a significant degree of overlap between corresponding images and text descriptions, affirming the rationale behind contrastive learning: aligning two modalities in the learned feature space. This alignment allowed the image and text encoders to mutually enhance each other, resulting in improved distinctiveness in image and text features compared to Fig. 5(c).

In Fig. 5(e), we present the confusion matrix for pathCLIP in the gene recognition task. While most gene names were recognized with high accuracy, some failures persisted. We exemplified these failures in Fig. 5(e). The primary reasons for these failures included low-quality pathway figures and imperfect text localization results from the object detection model. Pathway figures with low resolutions posed challenges for gene name recognition, even for human observers. Additionally, imperfect object detection occasionally led to image slices with incomplete gene names or extraneous words, contributing to recognition failures.

### D. Relation Extraction by pathCLIP

To underscore the added value brought by contrastive learning to pathCLIP, we conducted a comprehensive evaluation of relation extraction from pathway figures using different image encoders within the pathCLIP framework. We employed the GRE dataset (as detailed in Table I) for this evaluation. Table IV presents a comparative analysis of the performance achieved by the original ResNet50 (pre-trained on ImageNet [18]), the ResNet50 model from our previous work, and the ResNet50 model from pathCLIP for gene-relation extraction tasks on the detected relation image slices. Because the three methods are all based on the results of object detection, we additionally show the performance indicators of relation extraction after passing pathways through object detection and pathCLIP.

**TABLE IV.** Relation extraction performance of pathCLIP on detected relation image slices

| Category | Model | Precision [%] | Recall [%] | F1-score [%] |
| --- | --- | --- | --- | --- |
| Inhibit | **pathCLIP** | **96.79** | **90.40** | **93.49** |
|  | Original ResNet50 | 49.23 | 50.00 | 49.61 |
|  | Fine-tuned ResNet50 | 96.98 | 64.20 | 77.26 |
| Activate | **pathCLIP** | **90.99** | **97.00** | **93.90** |
|  | Original ResNet50 | 49.21 | 48.44 | 48.82 |
|  | Fine-tuned ResNet50 | 73.24 | 98.00 | 83.83 |
| Average | **pathCLIP** | **93.89** | **93.70** | **93.69** |
|  | Original ResNet50 | 49.22 | 49.22 | 49.21 |
|  | Fine-tuned ResNet50 | 85.11 | 81.10 | 80.52 |

From the evaluation metrics in Table IV, it became evident that ResNet50 from pathCLIP consistently outperformed both the original ResNet50 and the fine-tuned ResNet50 from our previous work in terms of precision, recall, and F1-score [34]. Notably, the evaluation metrics for the pre-trained ResNet50 model in the “activate” and “inhibit” categories hovered around 50%, indicating that without specialized fine-tuning using relevant images and labels, the model struggled to effectively classify these two categories. The superior performance of ResNet50 from pathCLIP revealed the benefits of the contrastive learning strategy, which harnessed additional guidance from textual modalities alongside image data.

Upon comparing the performance between the “activate” and “inhibit” categories, a distinct imbalance in our previous ResNet50 model becomes apparent. Specifically, the “inhibit” category achieved a precision score of 96.98% but a recall of 64.20%, whereas the “activate” category displayed a recall score of 98.00% but a precision of 73.24%. These results showed the relatively limited discriminative power of our previous ResNet50 model compared to ResNet50 from pathCLIP. The notable improvement exhibited by pathCLIP, with all scores exceeding 90% in both “activate” and “inhibit” categories, further validated the enhancement of the image encoder through support from the textual modality.

Furthermore, to emphasize the distinctiveness of embeddings generated by pathCLIP, we plotted the distribution of image encoders using UMAP dimension reduction, as shown in Fig. 6(b). Fig. 6(b) and Fig. 6(c) represented the distribution diagrams of image embeddings from the original ResNet50 and ResNet50 from pathCLIP, respectively. In Fig. 6(a), data points from the “activate” and “inhibit” categories overlap significantly, making differentiation challenging. However, in Fig. 6(c), the distribution of the two categories was obviously separate, aligning with their superior performance and validating the improvements brought by pathCLIP. Fig. 6(d) further illustrated the distributions of text features and image features from pathCLIP. Comparing Fig. 6(b) and Fig. 6(c), we observed that data points from the two categories in the embedding space shifted in disparate directions. This shift correlates with the positions of “inhibit text” and “activate text” data points in Fig. 6(d), presenting how guidance from text features steered the image embedding process, ultimately leading to attain distinctive qualities.

Finally, Fig. 6(e) showed the confusion matrix for the pathCLIP model in relation extraction. Promisingly, true positives and true negatives are prominent. However, we also found some instances of failure, including low-quality relation snippets and ambiguous samples. These challenges arose when relation snippets contained multiple gene relations due to crowded layouts in full pathway figures or when they only partially covered gene mentions due to imperfect object detection results. In some pathway figures, authors may depict reverse biological processes, including both “activate” and “inhibit” relations for a specific gene pair, leading to ambiguous cases and potential failures.

## IV. Case study

In a comprehensive analysis of 217 articles from PubMed specifically focusing on non-small cell lung cancer (NSCLC), we successfully extracted 612 genes and 1310 relations. These were subsequently imported into the Neo4j graph database, a robust platform tailored for intricate analyses and advanced research endeavors.

Fig. 7(a) delineates the elucidated pathway between EGFR and ERK as represented within Neo4j. Within this visualization, the green edges demarcate inhibition relations, whereas the red edges signify activation relations. Importantly, each edge is annotated with the PMID, referencing the originating source article for the specified relation. This portrayal unambiguously highlights several pivotal genes associated with NSCLC, notably including members of the RAS family, as well as AKT, PI3K, RAF, and MEK [26]. Crucially, the presented pathway wherein EGFR sequentially activates RAS, RAF, MEK, and ERK is in alignment with our current understanding of the gene pathway interconnecting EGFR and ERK.

**Fig. 7.**
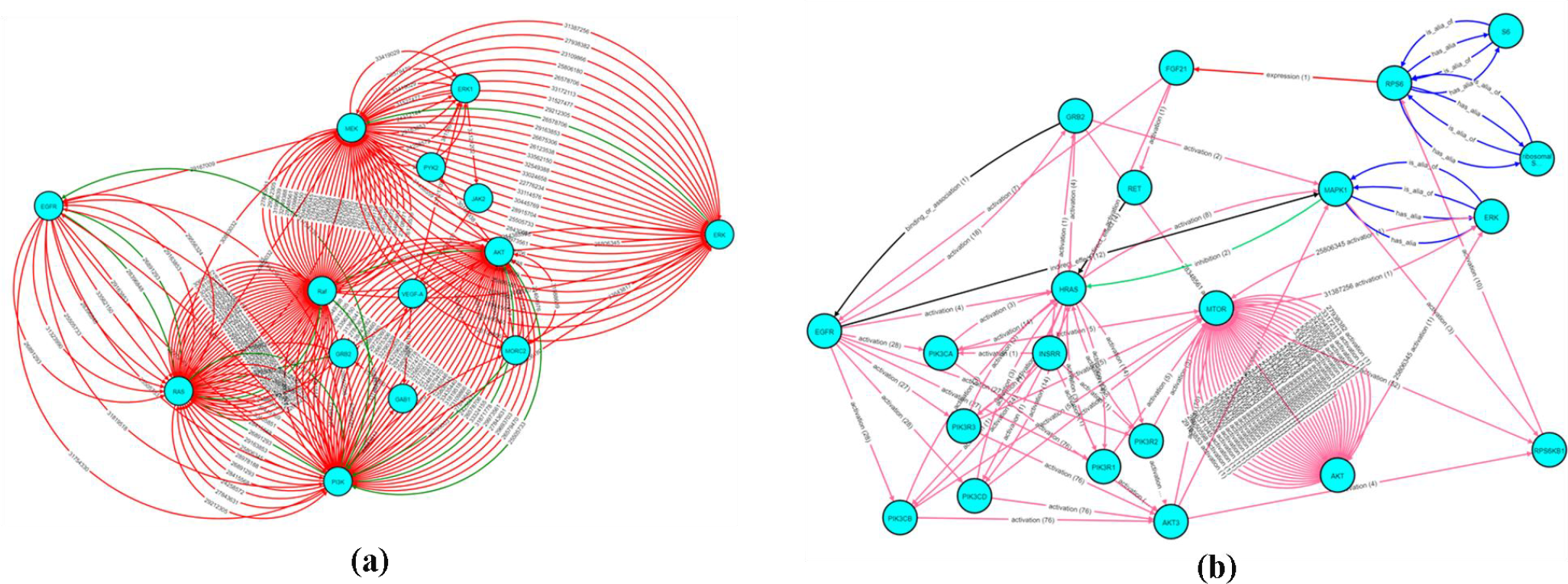
(a): Pathway between EGFR and ERK retrieved from Neo4j. (b)pathway between EGFR and ERK after enriching our database with KEGG and HUGO.

To augment the depth and breadth of our repository, we integrated the KEGG Pathway database, leveraging the KEGG API and the KEGG Markup Language (KGML) specifically designed for pathway maps. Furthermore, to encapsulate a broader gene and alias spectrum, we amalgamated the HUGO database (Braschi et al., 2019), achieving this by correlating HUGO ontology with the KEGG ontology.

Fig. 7(b) delineates the enhanced pathway between EGFR and ERK post integration of KEGG and HUGO data. Here, each edge represents a specific gene relation, classified either as activation, inhibition, alias, binding, or association. Furthermore, we have underscored each edge with its credibility measure, derived from the volume of supporting pathway maps or articles. It’s noteworthy to highlight that, in this enriched visualization, pivotal genes like PIK3CA, AKT, and HRAS remain evident. Additionally, the credibility of relations between two genes is augmented by cross-validation against the KEGG Pathway database. Notwithstanding the introduction of additional genes, it is evident that the core pathway between EGFR and ERK remains predominant, with the newer genes primarily serving as branches or ancillary pathways. Such precision underscores the value proposition of our methodology, enabling researchers to concentrate on highly pertinent gene pathways, thereby minimizing potential distractions.

A comparative analysis of Fig. 7 reveals nuances and contradictions, such as the divergent roles of “PI3K activates EGFR” versus “PI3K inhibits EGFR “. Such discrepancies may be indicative of methodological disparities or sample variations across different studies. Hence, our innovative approach of gleaning pathway insights directly from the scientific literature offers a robust mechanism to validate and enhance existing pathway databases. This, we believe, is an indispensable asset, pushing the frontiers of our understanding and potentially catalyzing advances in personalized medicine.

## V. Conclusion

In this paper, we introduced pathCLIP, a novel pathway curation system that seamlessly integrates image and text modalities to identify genes and gene relations within pathway figures. Our approach leveraged a contrastive learning framework to boost the image encoder with guidance from textual information, resulting in notable advancements in gene name recognition and gene-gene relation extraction from pathway figures. Through a series of comparative experiments and illuminating visualizations, we showcased the potency of contrastive learning, enhancing the discriminative capabilities of image embeddings for both gene recognition and gene-gene relation extraction tasks. These achievements compelled performance gains in pathway figure curation. While the challenges such as imperfect object detection and the presence of low-quality pathway figures leading to instances of pathway information extraction failure remained for our future works, our curations across multiple literature sources can enrich existing pathway databases and create a more comprehensive pathway map, facilitating advancements in biological research and precision medicine practices. These assertions were further validated through our case study on a newly collected pathway dataset derived from non-small cell lung cancer literature.

## Acknowledgment

We would like to acknowledge the valuable contribution from OpenAI’s GPT-3.5 model for aiding in language editing.

**Fei He** is a PhD student at the Department of Electrical Engineer and Computer Science of the University of Missouri. His research interests include bioinformatics applications of deep learning, single cell data analysis and protein structural analysis.

**Kai Liu** entered Northeast Normal University in 2021 to pursue his master’s degree, focusing on solving problems related to computer vision and computational biology using deep learning.

**Zhiyuan Yang** entered Northeast Normal University in China in 2022 to pursue her master’s degree. She is now dedicated to research in deep learning and bioinformatics. She has been involved in projects in the fields of image analysis and bioinformatics.

**Yibo Chen** is a Bioinformatics PhD candidate at the University of Missouri. He specializes in machine learning, bioinformatics, and software development, with a focus on Neo4j graph databases and programming in Python and Java. His research encompasses knowledge graphs, and biomedical text mining.

**Richard Hammer** is Professor and Vice Chair of Clinical Affairs, Pathology and Anatomical Sciences at University of Missouri. He focuses on providing state-of-the-art diagnosis and evaluation using the latest evidence-based medicine. He also is involved in bioinformatics and developing tools to apply digital solutions to clinical practice and clinical decision support.

**Mihail Popescu** is a full professor at University of Missouri-Columbia and works on biomedical intelligent systems and eldercare. His current research focuses on developing decision support systems for eldercare and chronic disease management. Chronic diseases such as diabetes, heart failure or depression decrease the quality of life and increase the risk of re-hospitalization. Dr. Popescu is currently investigating new technologies for eldercare such as new sensors, new monitoring technologies and new machine learning algorithms used to avoid this decrease. His research in early recognition of chronic conditions is intended to improve the quality of life and reduce the cost of health care. Dr. Popescu is developing novel summarization and visualization methodologies to address the health care big data problem associated with the patient activity monitoring process.

**Dong Xu** is Curators’ Distinguished Professor in the Department of Electrical Engineering and Computer Science, with appointments in the Christopher S. Bond Life Sciences Center and the Informatics Institute at the University of Missouri-Columbia. He obtained his Ph.D. from the University of Illinois, Urbana-Champaign in 1995 and did two years of postdoctoral work at the US National Cancer Institute. He was a Staff Scientist at Oak Ridge National Laboratory until 2003 before joining the University of Missouri, where he served as Department Chair of Computer Science during 2007-2016 and Director of Information Technology Program during 2017-2020. Over the past 30+ years, he has conducted research in many areas of computational biology and bioinformatics. His research since 2012 has focused on the interface between bioinformatics and deep learning. He was elected to the rank of American Association for the Advancement of Science (AAAS) Fellow in 2015 and American Institute for Medical and Biological Engineering (AIMBE) Fellow in 2020.

## Notes

This work was supported by the National Library of Medicine of the National Institute of Health (NIH) award number 5R01LM013392. The content is solely the responsibility of the authors and does not necessarily represent the official views of the National Institutes of Health. This work was also supported by the Science and Technology Development Program of Jilin Province of China (Grant No. 20210101174JC) to Kai Liu and Zhiyuan Yang.

### Competing Interest Statement

The authors have declared no competing interest.

